# Developmental changes within the genetic architecture of social communication behaviour: A multivariate study of genetic variance in unrelated individuals

**DOI:** 10.1101/179978

**Authors:** Beate St Pourcain, Lindon J Eaves, Susan M Ring, Simon E Fisher, Sarah Medland, David M Evans, George Davey Smith

**Affiliations:** Language and Genetics Department, Max Planck Institute for Psycholinguistics, The Netherlands; MRC Integrative Epidemiology Unit (MRC IEU), University of Bristol, UK; Donders Institute for Brain, Cognition and Behaviour, Radboud University, The Netherlands; Department of Human and Molecular Genetics, Institute for Psychiatric and Behavioral Genetics, Commonwealth University School of Medicine, Richmond, Virginia, USA; School of Social and Community Medicine, University of Bristol, UK; Psychiatric Genetics, QIMR Berghofer Medical Research Institute, Queensland, Australia; University of Queensland Diamantina Institute, Translational Research Institute, University of Queensland, Queensland, Australia

**Keywords:** ALSPAC, Structural equation modelling, Longitudinal analysis, Genetic variance decomposition, Genetic-relationship matrix structural equation modelling, Genetic relationship matrix

## Abstract

**Background:** Recent analyses of trait-disorder overlap suggest that psychiatric dimensions may relate to distinct sets of genes that exert their maximum influence during different periods of development. This includes analyses of social-communciation difficulties that share, depending on their developmental stage, stronger genetic links with either Autism Spectrum Disorder or schizophrenia. Here we developed a multivariate analysis framework in unrelated individuals to model directly the developmental profile of genetic influences contributing to complex traits, such as social-communication difficulties, during a ∼10-year period spanning childhood and adolescence.

**Methods:** Longitudinally assessed quantitative social-communication problems (N≤ 5,551) were studied in participants from a UK birth cohort (ALSPAC, 8 to 17 years). Using standardised measures, genetic architectures were investigated with novel multivariate genetic-relationship-matrix structural equation models (GSEM) incorporating whole-genome genotyping information. Analogous to twin research, GSEM included Cholesky decomposition, common pathway and independent pathway models.

**Results:** A 2-factor Cholesky decomposition model described the data best. One genetic factor was common to SCDC measures across development, the other accounted for independent variation at 11 years and later, consistent with distinct developmental profiles in trait-disorder overlap. Importantly, genetic factors operating at 8 years explained only ∼50% of the genetic variation at 17 years.

**Conclusion:** Using latent factor models, we identified developmental changes in the genetic architecture of social-communication difficulties that enhance the understanding of ASD and schizophrenia-related dimensions. More generally, GSEM present a framework for modelling shared genetic aetiologies between phenotypes and can provide prior information with respect to patterns and continuity of trait-disorder overlap.

## Introduction

The extent to which genetic aetiologies are shared between traits and disorders naturally depends on the genetic composition of the two phenotypes. While psychiatric disorders are diagnostic entities, defined by clinical criteria including the age of onset, human behaviour changes continously during development. This includes developmental alterations in compex genetic trait architectures as reported for cognitive (1) but also social-communication related characteristics (2).

Difficulties to socially engage and communicate with others, as observed in the general population, are heritable (twin-h^2^=0.74) (3) and a considerable proportion of the underlying genetic variation can be tagged by Single Nucleotide Polymorphisms (SNPs, SNP-h^2^≤ 0.45) (2). For both, social-communication and social interaction problems, multivariate twin (4;5) and bivariate GREML (genetic-relationship-matrix residual maximum likelihood) studies (6) reported evidence for a degree of genetic stability, but also change during childhood and adolescence (2;7;8) that may affect genetic similarities with other traits.

Studying the genetic overlap between psychatric illness and social-communciation difficulties across multiple developmental stages, different developmental profiles for childhood- versus adult-onset psychiatric disorders have been identified (9). The genetic overlap with clinical Autism Spectrum Disorder (ASD), a complex highly heritable early-onset neurodevelopmental condition (10), was strongest for social-communication difficulties during childhood, but declined with progressing age of the trait. By contrast, the genetic correlation with clinical schizophrenia, an adult-onset psychiatric illness with a typical first-time diagnosis between 16 to 30 years (10), was highest for social-communication problems during later adolescence (9). Thus, the risk of developing these contrasting psychiatric conditions might be related to distinct sets of genes, both of which affect social communication skills, but exert their maximum influence during different periods of development.

Discontinuity in trait-disorder overlap may, however, also result because of attrition-related artefacts such as decreasing power or inherent sample bias (11). As knowledge about developmental changes in complex genetic trait architectures is still scarce, development-related variations in trait-disorder overlap are often dismissed.

The aim of this work is to provide insight into the developmental profile of genetic factors influencing complex traits, such as social-communication difficulties during childhood and adolescence, using a longitudinal analysis framework. Building on our previous work (2;9), we investigate here two extreme hypotheses: We evaluate whether the genetic variance/covariance structure of social-communication difficulties during childhood and adolescence is consistent with multiple independent genetic influences, suggesting developmental changes in the genes responsible for inter-individual variation over time, or whether, alternatively, there is evidence for a shared single genetic factor, irrespective of age.

To study the developmental profile of genetic factors in unrelated individuals, we implemented multivariate genetic-relationship-matrix structural equation models (GSEM). These models utilise genome-wide genetic relationship matrices (GRMs)(12), calculated from hundreds of thousands of SNPs across the genome, to estimate the total amount of phenotypic variance and covariance tagged by common genetic variants, similar to GREML (12;13). GREML and related approaches (12;14–16) have re-shaped the research of complex genetic trait architectures beyond twin designs by exploiting the availability of genome-wide genetic data in cohorts of unrelated individuals. Genetic correlations are, however, typically estimated by these methods by studying two phenotypes only. Using a structural equation modelling (SEM) framework (17), as widely applied within twin research (4;5), we now extend this bivariate approach by flexibly modelling complex latent genetic factor structures within a multivariate context.

In this paper we use multivariate GSEM to model longitudinal data on social-communication difficulties across childhood and adolescence in the Avon Longitudinal Study of Parents and Children (ALSPAC), a phenotypically-rich longitudinal population-based birth cohort from the UK (18).

## Methods

### Participants and measures

All analyses were carried out using children’s data from ALSPAC, a UK population-based longitudinal pregnancy-ascertained birth-cohort (estimated birth date: 1991 to 1992)(18). Please note that the study website contains details of all the data that is available through a fully searchable data dictionary (http://www.bris.ac.uk/alspac/researchers/data-access/data-dictionary/). Ethical approval was obtained from the ALSPAC Law-and-Ethics Committee (IRB00003312) and the Local Research-Ethics Committees. Written informed consent was obtained from a parent or individual with parental responsibility and assent (and for older children consent) was obtained from the child participants.

#### Phenotype information

Social-communication difficulties during childhood and adolescence were collected with the 12-item mother-reported Social Communication Disorder Checklist (SCDC; score-range: 0 to 24, age range: 3 to 18 years)(3). The SCDC is a brief screening instrument of social reciprocity and verbal/nonverbal communication (e.g. "Not aware of other people’s feelings”), which has high reliability and internal consistency, and good validity (3) with higher scores reflecting more social-communication deficits. Quantitative SCDC scores in ALSPAC children and adolescents were repeatedly measured at 8, 11, 14 and 17 years and information on phenotypic and genotypic data was available for 4,174 to 5,551 children (Supplementary Table S1).

Descriptive analyses of SCDC scores were carried out in R.v.3.2.4. The distribution of SCDC scores was positively skewed and predominantly leptokurtic (Supplementary Table S1). Each score was adjusted for sex, age and the two most significant ancestry-informative principal components (see below) using ordinary least square (OLS) regression. Residuals were subsequently transformed to perfect normality using rank-based inverse normal transformation (19), as previously reported (9), to allow for comparisons across different algorithms (see below). There were moderate phenotypic correlations between repeatedly assessed SCDC scores, using both untransformed and transformed data (Supplementary Table S2, SCDC: Spearman’s-ρ: 0.39 to 0.57; Pearson-r: 0.38 to 0.61) as previously shown (9).

#### Genome-wide genotype information

ALSPAC children were genotyped using the Illumina HumanHap550 quad chip genotyping platforms (Supplementary Methods). After quality control, 8,237 children and 477,482 directly genotyped Single Nucleotide Polymorphisms (SNPs) were kept within the study.

### GSEM

Multivariate SEM techniques were used to assess the relative importance of genetic and residual influences to variation in longitudinal SCDC scores during child and adolescent development. Similar to GREML (12), GSEM use the genetic similarity between unrelated individuals to partition the expected phenotypic variance/covariance matrix into genetic and residual components. More generally, however, the statistical framework of GSEM is analogous to twin analysis methodologies (4;5), but uses GRMs, instead of twin correlations, to estimate genetic variance/covariance structures using full information maximum likelihood (FIML). Thus, genetic and environmental influences are modelled in the GSEM framework as latent factors contributing to inter-individual covariation in phenotypic measures. The advantage of our approach is that multivariate SEM methodology has been widely established within twin research (4;5) and allows for flexible modelling of complex genetic factor structures. Conversely, GREML, as implemented in the GCTA software package, is currently restricted to bivariate situations (20). While multivariate GSEM can be fit with SEM software such as OpenMx (21) using both mxGREML and FIML algorithms, these models are currently computationally expensive (see Results). We therefore implemented GSEM within R (Rv3.2.4) (for details see Supplementary Methods).

In short, GSEM describe the phenotypic covariance structure using one or more additive genetic factors A that capture genetic variance, tagged by common genotyped SNPs, as well as one or more residual factors E that capture residual variance, containing untagged genetic variation and unique environmental influences (including measurement error). As SEM methodology has its origins in the method of path analysis (22), path diagrams are useful in visualising the relationship among observed and latent variables (represented as squares and circles respectively, see e.g. Figure 2). Single headed arrows (factor loadings or ‘paths’) denote causal relationships between measures, whereas double headed arrows define correlations.

In our formulation, additive genetic variances (GSEM-Var_g_) and genetic covariances (GSEM-Cov_g_) are modelled as the product of additive genetic factor loadings and genetic factor variances (the latter being standardised to unit variance). For example, using multivariate GSEM, a saturated model can be fit to the data through a decomposition of both the genetic variance and residual variance into as many latent factors as there are observed variables (Cholesky decomposition model; see Supplemental methods). Estimated genetic variances and covariances can then be used to estimate genetic correlations (GSEM-r_g_) (23), the extent to which two phenotypes share common genetic factors (Supplementary Methods). Here, we utilised the Cholesky decomposition model as saturated and baseline model (Supplementary Information). Beside Cholesky decomposition models, multivariate GSEM also permit the fitting of models with smaller numbers of latent genetic and residual factors, defined according to theory (24).

Multivariate GSEM of longitudinally assessed SCDC scores were fitted in two stages.

In a first step (I), we specified *a priori* three standard multivariate AE models, analogous to twin research (Figure 2A-C): we studied a Cholesky decomposition model (saturated model), an independent pathway model and a common pathway model.

1. The Cholesky decomposition model, as described above, is a fully parametrised descriptive model without any restrictions on the structure of latent genetic and residual influences (20 free parameters) (Figure 2A) and involves multiple independent genetic influences sharing genetic aetiologies across development.
2. The independent pathway model, in its simplest form, specifies a single common genetic factor and a single common residual factor, in addition to age-specific genetic and residual influences (16 free parameters) (Figure 2B).
3. The common pathway model, in its simplest form, parametrises a single latent factor, influenced by both genetic and residual sources of variance, in addition to age-specific genetic and residual influences (Figure 2C), and is the most constrained model (14 free parameters). The model constrains the variance of the latent factor to one (i.e. the sum of squared genetic and residual factor loadings). Although the likelihood of this model can be estimated, the resulting Hessian is not invertible due to singularity problems. For these reasons, the model constraint was relaxed within this work.

Both, the independent pathway model and the common pathway model are consistent with a shared single genetic factor across development and are nested submodels of the full Cholesky decomposition model.

The goodness-of-fit of GSEM to empirical data was assessed using likelihood ratio test (LRT), the Akaike Information Criterion (AIC) (25) and the Bayesian Information Criterion (BIC) (26) (Supplementary Methods).

In a second step (II), we adopted a data-driven approach and investigated the pattern of genetic factor loadings for the best fitting model from (I) in detail. The smallest genetic factor loadings were successively dropped from the model and the overall fit of the model compared with the best-fitting *a priori* defined GSEM (or an adapted form) using LRTs. The statistical significance of factor loadings was assessed using a Wald test (2-sided test). Standard errors (SEs) for genetic and residual variances and covariances, and genetic correlations were derived from the variance-covariance matrix of the estimated factor loadings using the delta method. Standard errors for factor loadings were estimated by GSEM. Note that for rank-transformed measures with unit variance, such as the SCDC scores in this study, genetic variances are equivalent to SNP-h^2^ estimates. However, path coefficients for multivariate GSEM were re-standardised to enhance the interpretability.

GRMs were estimated using the GCTA software (12) and based on directly genotyped SNPs. All GSEM were fitted to data from participants with non-missing information to simplify the estimation algorithm. All R scripts are available via the R gsem package. (https://gitlab.gwdg.de/beate.stpourcain/gsem, Supplementary Information).

For the purpose of benchmark comparisons with univariate GCTA, we also fitted univariate GSEM, where genetic variances were estimated as a single variance component.

### GREML

The GCTA software package can be used to estimate the proportion of phenotypic variation that is jointly explained by SNPs on a genotyping chip using GREML (13) (AE model). Likewise, bivariate GREML (20) allows estimating genetic covariances and genetic correlations between two phenotypes. An advantage of this method is that genetic correlations between two phenotypes can be estimated even when these phenotypes are not measured in the same individuals.

Univariate and bivariate GREML were carried out as part of sensitivity and simulation analyses. For comparison with GSEM, genetic relationship matrices (GRMs) were derived from directly genotyped SNPs, but excluded individuals with a pairwise relationship >0.025, as recommended (13). All analyses were conducted with GCTA software v1.25.2 (12).

#### OpenMx SEM models

OpenMx SEM models (21), as implemented in the OpenMx software (http://openmx.psyc.virginia.edu/)(v2.5 and v2.7), were fitted using FIML and mxGREML and included a full Cholesky decomposition of both genetic and residual variances (AE model, see above). Bivariate OpenMx SEM analyses were conducted as part of a simulation analysis. Genetic variances, genetic covariances, and genetic correlations were derived as described for GSEM above.

All analyses were conducted on High Performance Clusters at the University of Bristol and the MPI for Psycholinguistics.

#### Data simulation

To evaluate the accuracy of multivariate GSEM, we carried out data simulations (Supplementary Methods).

#### Attrition analysis

SCDC-attrition scores were generated to investigate potential sources of bias. Analyses included sample-specific estimates of genetic correlations among SCDC-attrition scores, and between SCDC scores and subsequent sample dropout (Supplementary Methods).

## Results

### Accuracy of multivariate GSEM

We simulated a bivariate trait (N=5000) with two standardised measures (10 replicates; Supplementary Figure S1A, Supplementary Table S3) and confirmed the accuracy of multivariate GSEM through comparison with GCTA and OpenMx software. All methods provided accurate estimates, both with respect to genetic and residual variances and covariances as well as genetic and residual factor loadings (GSEM and OpenMx SEM models only), with comparable RMSE, MAD and little bias (Bias^2^<10^−3^ for all methods, Supplementary Table S3). Computationally, multivariate OpenMx SEM models were, however, more expensive (≤ 78 GB RAM FIML v2.5; ≤ 2694 minutes mxGREML/FIML v2.7) than multivariate GSEM (≤ 13 GB RAM, ≤ 301 minutes) per single bivariate replicate analysis. A comparison of computing resources is shown in Supplementary Table S4. There was also little difference between estimated OpenMx versus GSEM parameters when analysing a trivariate simulated trait with three standardised measures, as part of a benchmark test (Supplementary Figure S1B, Supplementary Table S5). Note that trivariate replicate analyses using OpenMx were not considered within this study due to computational constraints.

### Univariate analyses

Using univariate GSEM, common genetic variants explained a large proportion of phenotypic variation in SCDC scores during childhood as well as during later adolescence (age 8: Var_g_(SE)=0.25(0.061), *p*=3.4x10^−5^; age 11: Var_g_(SE)=0.22(0.061), *p*=2.9x10^−4^, age 17: Var_g_(SE)=0.47(0.086), *p*=4.4x10^−8^; Figure 1, Supplementary Table S6) but not during early adolescence (age 14, Var_g_(SE)= 0.086(0.064), *p*=0.18), as previously reported (2). Univariate GCTA(GREML) yielded nearly identical results (Supplementary Table S7).

**Figure 1:**
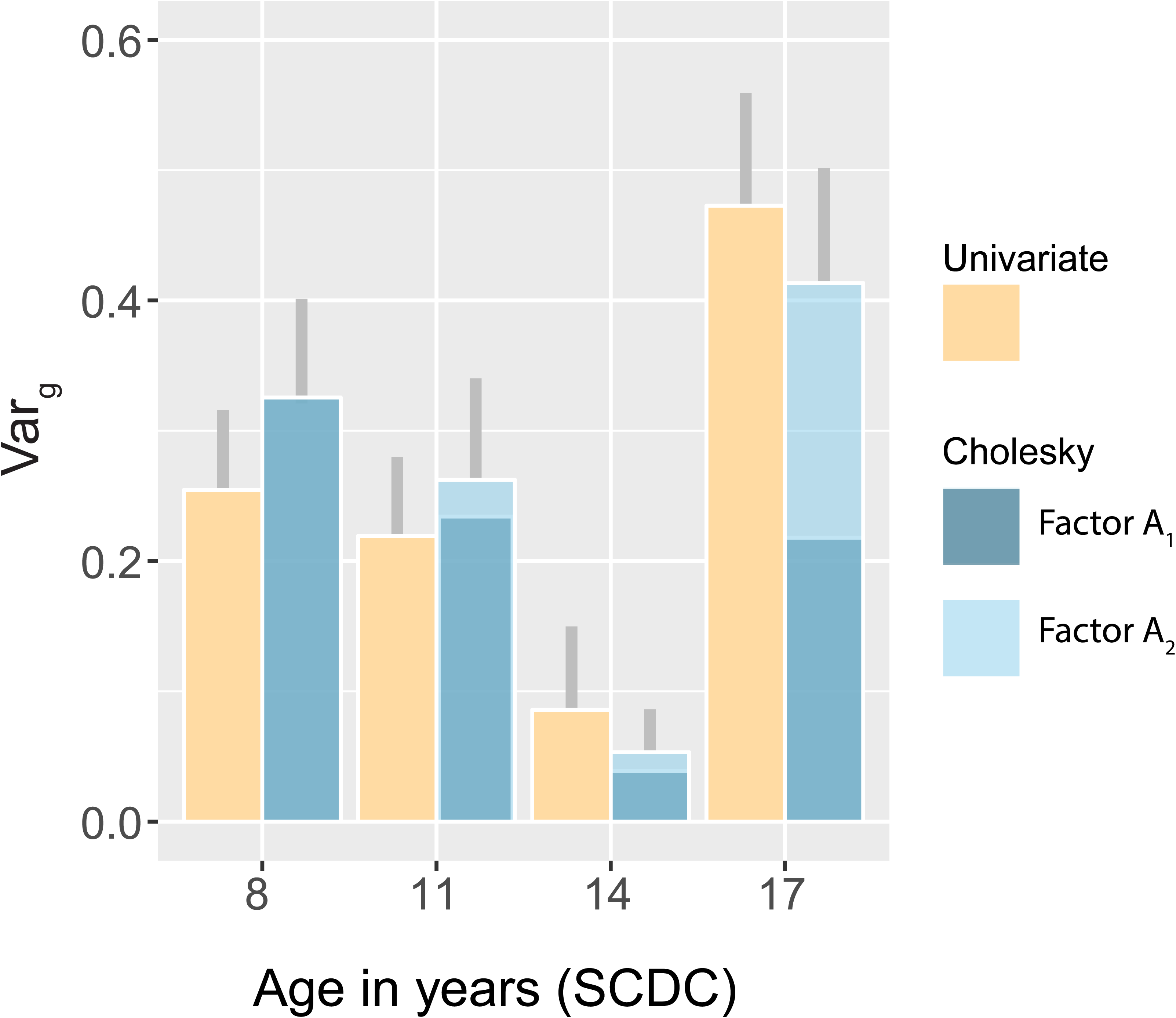
Genetic variance of SCDC scores during development. Genetic variances for SCDC scores across development as estimated using a univariate model (Supplementary Table S6, N≥ 4,174) and the full Cholesky decomposition model (Table 1, Model 1; Supplementary Table S8, N=3,295). Genetic factor A_3_ and A_4_ of the Cholesky decomposition model are not shown as their estimated Var_g_ was negligible (<0.01). All reported Var_g_ estimates are equivalent to SNP-h^2^ estimates. Grey lines indicate one standard error (SE) in total genetic variance (Var_g_) for each SCDC measure. SCDC-Social and Communication Disorders Checklist; Var_g_ - Genetic variance

#### Multivariate analyses

We first examined the profile of genetic factors contributing to variation in SCDC scores during development (13,180 observations; 3,295 participants) using three *a priori* defined multivariate GSEM (Figure 2A-C). Based on all three fit indices, LRT, AIC and BIC, the best-fitting *a priori* defined model was the full Cholesky decomposition model (Model 1, Table 1, Figure 2A, Figure 3A). Neither a single factor independent pathway model nor a single factor common pathway model could sufficiently capture the underlying variance/covariance structure of the data. As the full Cholesky decomposition model is, however, also the baseline model, the model identification progressed with the identification of meaningful GSEM through data-driven model modifications. Consistent with near zero factor loadings for the latent genetic factors A_3_ and A_4_ (Supplementary Table S8), a two genetic factor Cholesky model was studied (Model 4, Figure 2D) that provided a near-identical fit to the data (Table 1, ΔX^2^ <0.01 (Δdf=3), *p*=1). This model parametrised one genetic factor arising at age 8 years, and a second independent genetic factor explaining novel genetic influences arising at age 11 years, each contributing to phenotypic variation during later development (Figure 2D). Using LRTs, the model fitting progressed (Model 5, Table 1, Supplementary Table S8) until all genetic factor loadings reached *p*<0.05 without a significant drop in the log-likelihood (ΔX^2^=<0.01 (Δdf=2), *p*=1, with respect to Model 4).

**Figure 2:**
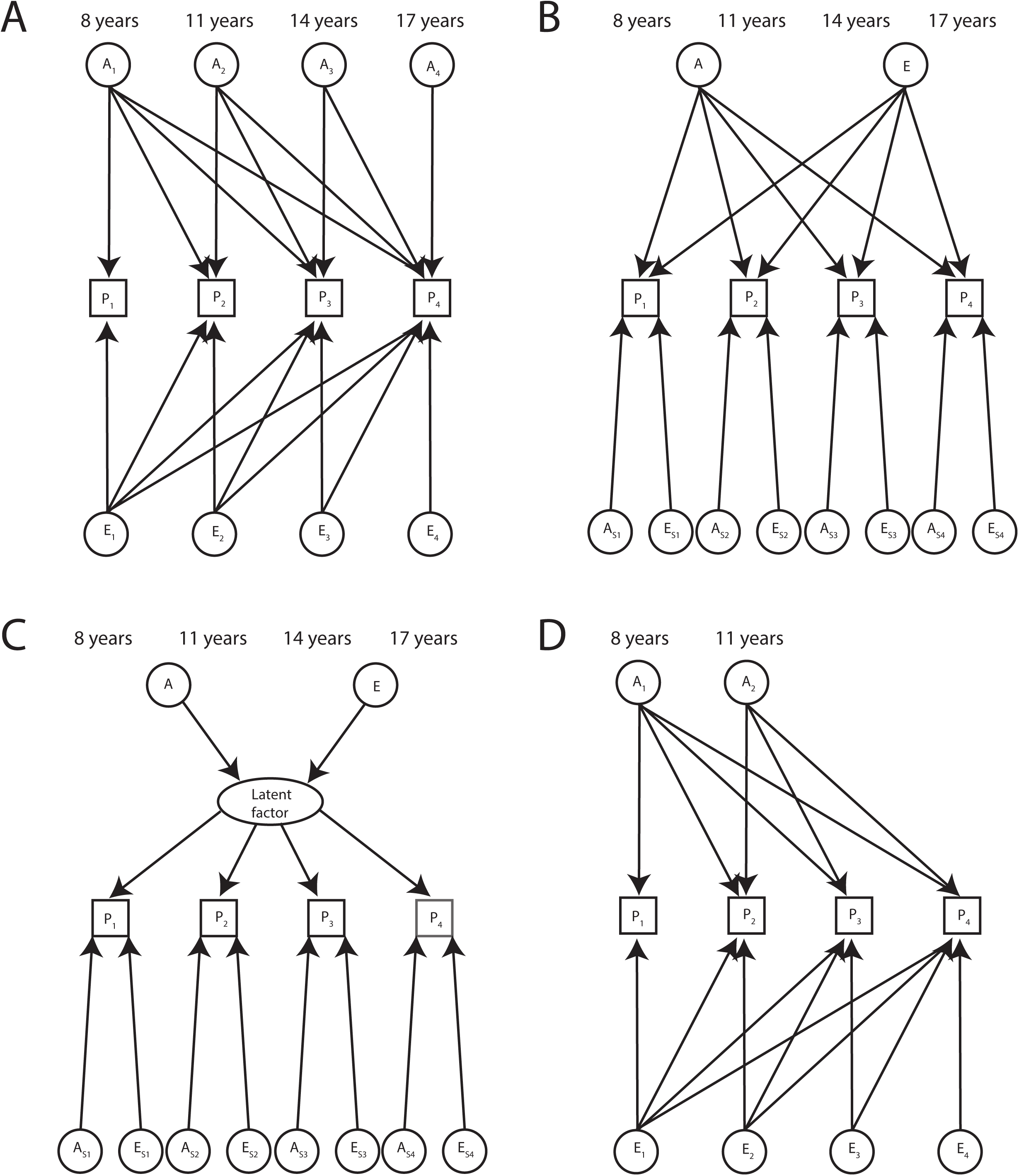
Path diagrams of *a priori* defined multivariate GSEM and data-driven model modifications. A - Full Cholesky decomposition model; B - Independent pathway model; C - Common pathway model; D - Two genetic factor Cholesky model (Data-driven model modification) Observed phenotypic measures are represented by squares and latent factors by circles. Single headed arrows (’paths’) define causal relationships between variables. Double headed arrows define correlations. Note that the variance of latent variables is constrained to unit variance, this is omitted from the diagrams to improve clarity. GSEM - Genetic-relationship-matrix structural equation models

**Table 1:** Multivariate GSEM of SCDC scores

| Model | Path diagram | -2LL | k | $\Delta X^2$ to model 1 | $\Delta df$ to model 1 | *p* | AIC | BIC |
| --- | --- | --- | --- | --- | --- | --- | --- | --- |
| ***A priori* defined multivariate GSEM** |  |  |  |  |  |  |  |  |
| 1. Full Cholesky decomposition model - *saturated model* | Figure 2A,<br>Figure 3A | 7900.97 | 20 | - | - | - | 7940.97 | 8062.97 |
| 2. Independent pathway model | Figure 2B | 7914.51 | 16 | 13.55 | 4 | 0.0089 | 7946.51 | 8044.12 |
| 3. Common pathway model | Figure 2C | 8082.7 | 14 | 181.73 | 6 | $<10^{-15}$ | 8110.70 | 8196.10 |
| **Data-driven model modification** |  |  |  |  |  |  |  |  |
| 4. Two genetic factor Cholesky model | Figure 2D | 7900.96 | 17 | $<0.01$ | 3 | 1 | 7934.96 | 8038.67 |
| | Path diagram | -2LL | k | $\Delta X^2$ to model 4 | $\Delta df$ to model 4 | *p* | AIC | BIC |
| **Best-fitting model** |  |  |  |  |  |  |  |  |
| 5. Two genetic factor Cholesky model (excluding non-significant paths) * | Figure 3B | 7900.96 | 15 | $<0.01$ | 2 | 1 | 7930.96 | 8022.47 |

**Figure 3:**
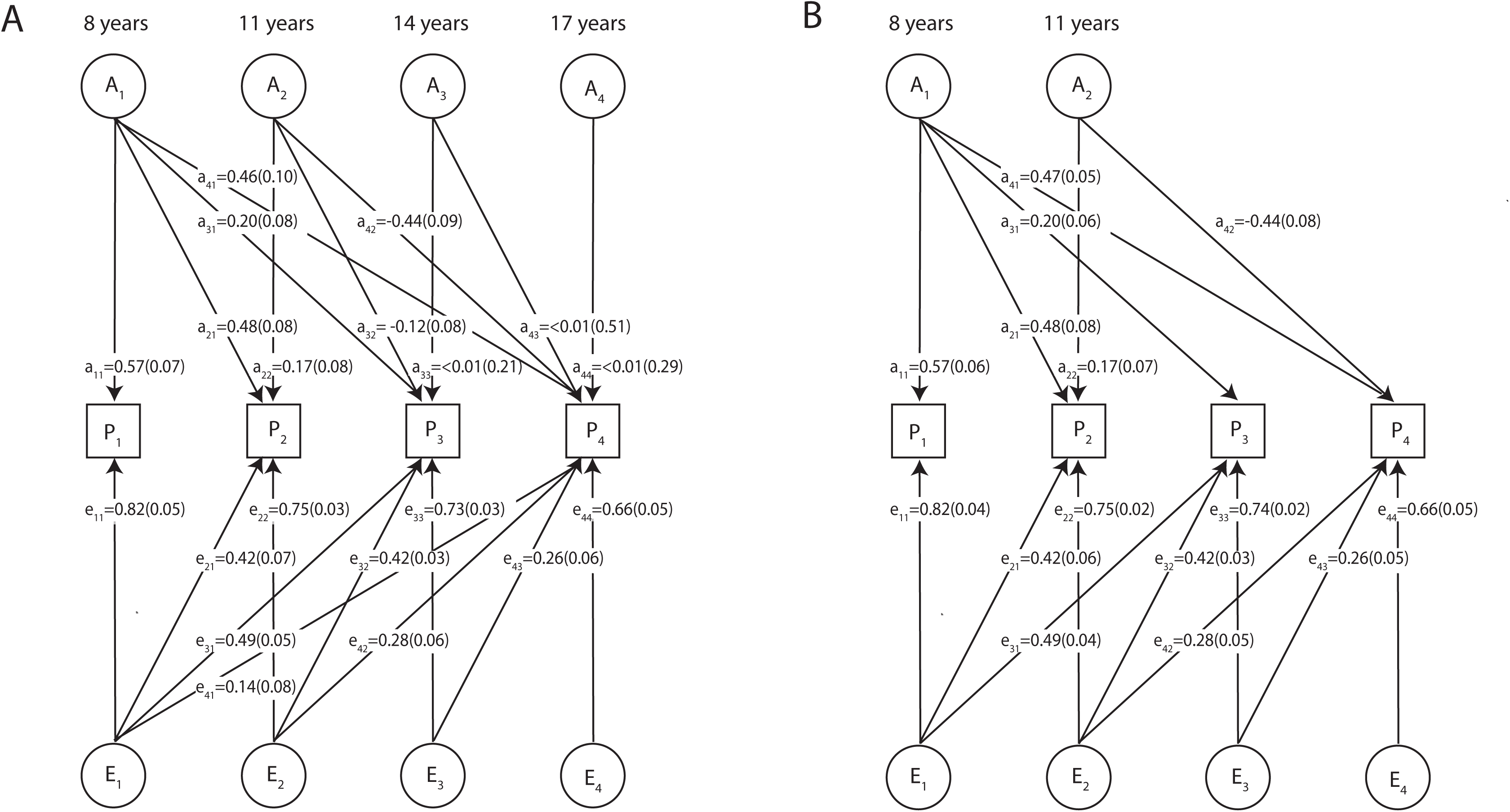
Path diagram of the full Cholesky decomposition model for SCDC scores (A) and its reduced form (B) The full Cholesky decomposition model (A) and its most parsimonious reduced form (B) are described in detail in Table 1 (Model 1 and 5 respectively). Corresponding to the phenotypic measures P_1_(8 years), P_2_(11 years), P_3_(14 years) and P_4_(17 years), the latent genetic factors with factor loadings a are A_1_(8 years), factor A_2_(11 years), factor A_3_(14 years), factor A_4_(17 years) and the latent residual factors with factor loadings e are E_1_(8 years), factor E_2_(11 years), factor E_3_(14 years), factor E_4_(17 years). All path coefficients are standardised. 3,295 participants had repeated scores across all ages. Note that the variance of latent variables is constrained to unit variance, this is omitted from the diagrams to improve clarity. SCDC - Social and Communication Disorders Checklist

The identified model included one common genetic factor A_1_, accounting for shared phenotypic variation throughout development, as well as a second genetic factor A_2_ influencing SCDC scores at 11 years and especially at 17 years of age (Table 1, Figure 3B). Figure 3 shows the full Cholesky decomposition model (Model 1) and its best-fitting reduced form (Model 5) with their standardised path coefficients (factor loadings ≥ 0.32 explain >10% of the phenotypic variance).

Overall, the estimates of genetic variance, as predicted by GSEM (Model 1 and 5, Supplementary Table S9), were consistent with univariate GSEM estimates (Figure 1), although latter were based on larger sample numbers (Supplementary Table S6). The pattern of genetic factor loadings suggested, however, a dynamic change in the variance composition of the trait during development such that only ∼50% of the genetic variance at age 17 was accounted for by genetic variation at age 8 (e.g. age 17: ratio Var_g_(A_1_) to Var_g_(A_1_+A2); Model 1: 0.53(SE=0.18)%; Model 5: 0.53(SE=0.12)%)(Figure 1).

The predicted bivariate genetic correlations by multivariate GSEM (Model 1 and 5, Supplementary Table S9) were overall similar to bivariate GCTA(GREML) estimates, although latter were based on larger numbers of observations (Supplementary Table S10 and Supplementary Figure S3). Restricting analyses to the same sets of individuals, both bivariate GSEM and bivariate GCTA(GREML) provided near-identical estimates (Supplementary Table S10), although these analyses were less powerful. Thus, small differences in genetic correlations patterns, as estimated by multivariate GSEM versus bivariate GCTA(GREML), are likely to be due to minor differences in sample numbers.

There was furthermore little evidence that genetic influences between SCDC scores and subsequent SCDC sample dropout are shared in ALSPAC (Supplementary Table S11). Nominal evidence for a genetic correlation was observed between SCDC scores at 8 years and dropout at 14 years only (r_g_=0.39(SE=0.19), *p*_*one-tailed*_=0.02). Nonetheless, SCDC attrition scores were genetically correlated across all SCDC measures in ALSPAC (*p*_*one-tailed*_<10^−3^, Supplementary Table S12).

### Discussion

Using multivariate SEM in combination with common variant-based genetic correlation matrices, we investigated the developmental structure of genetic factors contributing to social-communication difficulties during childhood and adolescence. We showed that the genetic architecture of this population-based complex trait changes continuously during development and is consistent with multiple genetic influences operating at different stages during development. Thus, our study provides evidence against the hypothesis that social communication behaviour during development is a genetically homogenous phenotype.

The best-fitting model, specifying two distinct genetic factors, suggested that the genetic origins of child and adolescent social-communication behaviour lie in middle and late childhood. The first genetic factor, parametrised to account for all genetic influences at age 8 years, explained a considerable proportion of phenotypic variance throughout development (>20%) with the exclusion of SCDC scores at age 14 that have negligible SNP-h^2^ estimates. (This is consistent with recent reports of low SNP-h^2^ for autistic symptoms at the beginning of adolescence (1) and might be related to pubertal adjustments (2)).

The second genetic factor, parametrised to be independent of the first one and to capture novel genetic influences arising at age 11 years, explained predominantly phenotypic variation at 17 years of age (∼19%). Thus, the model predicted changes in the composition of the genetic variance during development, and only ∼50% of the genetic variation at age 17 was accounted for by genetic variation at age 8. Within defined developmental stages, however, such as those spanning mid-childhood to very early adolescence (e.g. 8 to 11 years), we found evidence for strong genetic correlations across measures. These results are consistent with recent longitudinal twin research that reported moderate to high genetic stability for autistic traits, including communication impairments, between mid-childhood and early adolescence (7), but only moderate genetic stability between behaviour in childhood versus emerging adulthood (8). The identified genetic factor structure using GSEM reflects therefore both a degree of genetic stability, but also genetic change in social-communication behaviour during development, depending on the size of the developmental window.

The identification of two distinct genetic factors, especially during later adolescence, suggests that SCDC scores at age 8 or 11 years are, in terms of average composition, different from those influencing SCDC scores at age 17. Developmental changes in the genetic architecture of social communication traits are consistent with biological maturation processes during childhood and adolescence. For example, synaptic pruning in the cerebral cortex is a signature late maturational process for generating a diversity of neuronal connections (27), which occurs during puberty and extends into early adult life (28). In parallel, there are changes in adolescent social cognitive development, especially with respect to emotional perspective taking, resistance to peer influence and changes in social behaviour (29). Given the identified genetic factor structure, it could be speculated whether multiple concepts of ‘social reciprocity and verbal/nonverbal communication’ may co-exist, especially at age 17, and whether changes in genetic factor contributions may continue into early adulthood. Thus, even for psychological instruments with high reliability, internal consistency and good discriminant validity, like the SCDC (3), the nature of the captured continuous phenotype may vary across developmental periods spanning ∼10 years. This underlines the need for behavioural genetic studies across the life-span.

An important implication that flows from the observation of developmental variations in the genetic trait architecture is that measures assessed at different developmental stages may reveal different patterns of trait-disorder overlap, as previously shown for clinical ASD and schizophrenia respectively (9). Moreover, the identification of a 2-genetic factor is also consistent with recent reports of little genetic overlap between ASD versus schizophrenia-related dimensions (30), especially with respect to social-communication symptoms. Structural models capturing developmental changes in the genetic architecture of complex phenotypes can therefore be leveraged to obtain prior information concerning the stability of trait-disorder overlap and consequently the extent to which development-specific genetic trait factors are shared among different psychiatric dimensions.

Our findings have therefore specific relevance for the study of functional dimensions of human behaviour spanning the continua from normal to abnormal and across development, consistent with the framework of Research Domain Criteria (31).

Finally, our study proves that structural models of genetic influences in unrelated individuals, as captured by GRMs, are computationally feasible within a longitudinal context. Beyond the scope of bivariate GCTA(GREML), multivariate GSEM allow for the modelling of complex latent genetic factor structures across different stages of development, in particular their genetic variance composition, and can reveal developmental origins of genetic variation that are otherwise hidden. It is furthermore possible to envisage that the concept of GSEM can be extended to investigate multivariate models of cross-disorder overlap and other complex phenomena, such as reciprocal causation. Note that also novel OpenMx FIML and mxGREML algorithms are currently being developed.

A limitation of our study is the analysis of non-missing data across all repeatedly assessed measures. Thus, weaker genetic links, spanning wider age gaps, may not have been sufficiently captured as a consequence of lower power, although genetic correlations predicted by multivariate GSEM and bivariate GCTA(GREML) were overall similar. In addition, cohort studies can be affected by attrition bias (32). We identified, however, little evidence for a specific genetic link between variation in SCDC scores and subsequent sample dropout, although attrition scores across all assessed SCDC measures were genetically correlated. This is consistent with studies reporting an association between study non-participation, including SCDC dropout, and polygenic risk for schizophrenia (9;32), irrespective of when phenotypes were sampled during development. In addition, we exclusively studied rank-transformed phenotypes to ensure multivariate normality and comparability across different estimation algorithms, and we can therefore not exclude transformation-related biases. However, genetic overlap with psychiatric conditions provided some evidence for the content validity of the analysed trait (9). Also, maternal characteristics may have contributed to phenotypic and, to a lesser extent, genetic correlations. However, the impact of these effects is likely to be small, given the identified developmental changes in genetic variances and covariances for SCDC scores during development. Finally, a Cholesky decomposition of a variance/covariance matrix may not always result in fitting statistics that follow the expected chi-squared distribution (33). Model comparisons using real and simulated data provided, however, little evidence for systematic differences between GCTA(GREML), GSEM and OpenMx SEMs. Thus, despite potential limitations, our study demonstrates that structural models of longitudinally assessed behavioural traits can inform on developmental changes in genetic trait architectures as tagged by common SNPs.

### Conclusions

The genetic architecture of social-communication difficulties, as tagged by common genetic variation, changes with age and involves multiple genetic factors operating at different developmental stages during a 10-year period spanning childhood and adolescence. The identification of distinct genetic trait factors is consistent with different profiles of trait-disorder overlap, and underlines the importance of investigating genetic trait variances within a multivariate context.

## Acknowledgments

We are extremely grateful to all the families who took part in this study, the midwives for their help in recruiting them, and the whole ALSPAC team, which includes interviewers, computer and laboratory technicians, clerical workers, research scientists, volunteers, managers, receptionists and nurses. This publication is the work of the authors and they will serve as guarantors for the contents of this paper. The UK Medical Research Council and the Wellcome Trust (102215/2/13/2) and the University of Bristol provide core support for ALSPAC. The ALSPAC GWAS data was generated by Sample Logistics and Genotyping Facilities at the Wellcome Trust Sanger Institute and LabCorp (Laboratory Corporation of America) using financial support from 23andMe. Autism Speaks (7132) provided support for autistic-trait related analyses in ALSPAC to BSP. BSP and SF are supported by the Max Planck Society. We thank Robert Kirkpatrick for helpful discussions on structural equation models and support with the OpenMx code. We thank Gregory Carey for his contribution and many helpful discussions as part of the initial work on bivariate mGCTA models carried out together with LE, DE and BSP. We thank Callum Wright and Tobias van Valkenhoef for their help with the High Performance Computing systems.

## References

1. Trzaskowski M, Yang J, Visscher PM, Plomin R (2014): DNA evidence for strong genetic stability and increasing heritability of intelligence from age 7 to 12. Mol. Psychiatry 19: 380–384.

2. St Pourcain B, Skuse DH, Mandy WP, Wang K, Hakonarson H, Timpson NJ, et al. (2014): Variability in the common genetic architecture of social-communication spectrum phenotypes during childhood and adolescence. Mol. Autism 5: 18.

3. Skuse DH, Mandy WPL, Scourfield J (2005): Measuring autistic traits: heritability, reliability and validity of the Social and Communication Disorders Checklist. Br. J. Psychiatry 187: 568–572.

4. Neale M, Maes HHM (2004): Methodology for genetic studies of twins and families., Dordrecht: Kluwer Academic Publishers.

5. Martin NG, Eaves LJ (1977): The genetical analysis of covariance structure. Heredity 38: 79–95.

6. Lee SH, Yang J, Goddard ME, Visscher PM, Wray NR (2012): Estimation of pleiotropy between complex diseases using single-nucleotide polymorphism-derived genomic relationships and restricted maximum likelihood. Bioinformatics 28: 2540– 2542.

7. Holmboe K, Rijsdijk FV, Hallett V, Happé F, Plomin R, Ronald A (2014): Strong Genetic Influences on the Stability of Autistic Traits in Childhood. J. Am. Acad. Child Adolesc. Psychiatry 53: 221–230.

8. Taylor MJ, Gillberg C, Lichtenstein P, Lundström S (2017): Etiological influences on the stability of autistic traits from childhood to early adulthood: evidence from a twin study. Mol. Autism 8: 5.

9. St Pourcain B, Robinson EB, Anttila V, Bulik-Sullivan B, Maller JB, Golding J, et al. (2017): ASD and schizophrenia show distinct developmental profiles in common genetic overlap with population-based social-communication difficulties. Mol. Psychiatry Advance online publication: 10.1038/mp.2016.198.

10. American Psychiatric Association (1994): Diagnostic and Statistical Manual of Mental Disorders, 4th ed. Washington, DC: American Psychiatric Association.

11. Martin J, Tilling K, Hubbard L, Stergiakouli E, Thapar A, Smith GD, et al. (2016): Association of Genetic Risk for Schizophrenia With Nonparticipation Over Time in a Population-Based Cohort Study. Am. J. Epidemiol.10.1093/aje/kww009.

12. Yang J, Lee SH, Goddard ME, Visscher PM (2011): GCTA: A Tool for Genome-wide Complex Trait Analysis. Am. J. Hum. Genet. 88: 76–82.

13. Yang J, Benyamin B, McEvoy BP, Gordon S, Henders AK, Nyholt DR, et al. (2010): Common SNPs explain a large proportion of the heritability for human height. Nat. Genet. 42: 565–569.

14. Bulik-Sullivan BK, Loh P-R, Finucane HK, Ripke S, Yang J, Schizophrenia Working Group of the Psychiatric Genomics Consortium, et al. (2015): LD Score regression distinguishes confounding from polygenicity in genome-wide association studies. Nat. Genet. 47: 291–295.

15. Loh P-R, Tucker G, Bulik-Sullivan BK, Vilhjálmsson BJ, Finucane HK, Salem RM, et al. (2015): Efficient Bayesian mixed-model analysis increases association power in large cohorts. Nat. Genet. 47: 284–290.

16. Speed D, Hemani G, Johnson MR, Balding DJ (2012): Improved Heritability Estimation from Genome-wide SNPs. Am. J. Hum. Genet. 91: 1011–1021.

17. Bollen KA (1989): Structural Equations with Latent Variables, 1 edition. New York: Wiley-Blackwell.

18. Boyd A, Golding J, Macleod J, Lawlor DA, Fraser A, Henderson J, et al. (2013): Cohort Profile: The “Children of the 90s”—the Index Offspring of the Avon Longitudinal Study of Parents and Children. Int. J. Epidemiol. 42: 111–27.

19. Peng B, Yu RK, DeHoff KL, Amos CI (2007): Normalizing a large number of quantitative traits using empirical normal quantile transformation. BMC Proc. 1: S156.

20. Lee SH, Yang J, Goddard ME, Visscher PM, Wray NR (2012): Estimation of pleiotropy between complex diseases using single-nucleotide polymorphism-derived genomic relationships and restricted maximum likelihood. Bioinformatics 28: 2540– 2542.

21. Boker S, Neale M, Maes H, Wilde M, Spiegel M, Brick T, et al. (2011): OpenMx: An Open Source Extended Structural Equation Modeling Framework. Psychometrika 76: 306–317.

22. Wright S (1921): Correlation and causation. J. Agric. Res. 20: 557–585.

23. Falconer PDS, Mackay PTFC (1995): Introduction to Quantitative Genetics, 4 edition. Essex, England: Longman.

24. MacCallum RC, Roznowski M, Necowitz LB (1992): Model modifications in covariance structure analysis: The problem of capitalization on chance. Psychol. Bull. 111: 490–504.

25. Akaike H (1987): Factor analysis and AIC. Psychometrika 52: 317–332.

26. Schwarz G (1978): Estimating the Dimension of a Model. Ann. Stat. 6: 461–464.

27. Selemon LD (2013): A role for synaptic plasticity in the adolescent development of executive function. Transl. Psychiatry 3: e238.

28. Petanjek Z, Judaš M, Šimić G, Rašin MR, Uylings HBM, Rakic P, et al. (2011): Extraordinary neoteny of synaptic spines in the human prefrontal cortex. Proc. Natl. Acad. Sci. U. S. A. 108: 13281–13286.

29. Burnett S, Blakemore S-J (2009): The Development of Adolescent Social Cognition. Ann. N. Y. Acad. Sci. 1167: 51–56.

30. Taylor MJ, Robinson EB, Happé F, Bolton P, Freeman D, Ronald A (2015): A longitudinal twin study of the association between childhood autistic traits and psychotic experiences in adolescence. Mol. Autism 6:

31. Cuthbert BN, Insel TR (2013): Toward the future of psychiatric diagnosis: the seven pillars of RDoC. BMC Med. 11: 126.

32. Martin J, Tilling K, Hubbard L, Stergiakouli E, Thapar A, Smith GD, et al. (2016): Association of Genetic Risk for Schizophrenia With Nonparticipation Over Time in a Population-Based Cohort Study. Am. J. Epidemiol.kww009.

33. Carey G (2005): Cholesky Problems. Behav. Genet. 35: 653–665.

